# Attitudes and Practices of Open Data, Preprinting, and Peer-review - a Cross Sectional Study on Croatian Scientists

**DOI:** 10.1101/2020.11.25.395376

**Authors:** K Baždarić, I Vrkić, E Arh, M Mavrinac, M Gligora Marković, L Bilić-Zulle, J Stojanovski, M. Malički

## Abstract

Attitudes towards open peer review, open data and use of preprints influence scientists’ engagement with those practices. Yet there is a relatively small number of validated questionnaires that measure these attitudes. The goal of our study was to construct and validate such a questionnaire. Using a sample of Croatian scientists (N=541), from a wide range of disciplines, we developed a questionnaire titled Attitudes towards Open data sharing, preprinting, and peer-review (ATOPP). The questionnaire has 21 item with a four-factor structure (attitudes towards: open data, preprint servers, open peer-review and open peer-review in small scientific communities). Based on the questionnaire, the attitudes of Croatian scientists towards these topics were generally neutral, with a median of 3.3 out of 5 of the total attitude score. Croatian scientist attitudes were lowest for open peer-review in small scientific communities (Md 2.0) and highest for open data (Md 3.9).

## Introduction

Open science, despite lacking an universally accepted definition, is widely recognized as a global phenomenon and an initiative emerging from the philosophical concept of scholarly „openness“. With principles and values of openness rooted in the idea of scientific knowledge as a common good (1). The term open science was coined in 2001 by Recep Şentürk and he used it to refer to a democratic and a pluralist culture of science. For Şentürk, open science indicated that different perspectives in science are considered equal, rather than alternative to each other: “If we desire to recognize the complexity of our world we must embrace multiplex ontology” (1). His view, however, is different from today’s relatively narrow view of open science perceived as an „effort by researchers, governments, research funding agencies or the scientific community itself to make the primary outputs of publicly funded research results – publications and the research data – publicly accessible in digital format with no or minimal restriction“(2). A recent systematic review summarized definitions of open science from 75 studies into „transparent and accessible knowledge that is shared and developed through collaborative networks“(3).

Practical considerations of open science often dealt with methods to lower or erase technical, social, and cultural barriers, and enable public sharing of research plans and outputs (4), which are believed to lead toward the betterment of science (5). Often, those practical considerations are described in various open science taxonomies and classifications, of which one of the most commonly used - the FOSTER’s project graphical representation distinguishes six „first level” elements of open science: open access, open data, open reproducible research, open science evaluation, open science policies, and open science tools (6).

In our research, we focused on the three of these elements: open data, open science evaluation (open peer review), and open science tools (open repositories - preprint servers).

### Open peer-review

Open peer-review also lack an universal definition applied to the term (7). Most often it used to describe one of the following practices: open identities of the authors and reviewers, open review reports published alongside the article, open participation of the wider community, open interaction and discussion between author(s) and reviewers, open peer-review manuscripts which are made immediately available, open final-version commenting, and open platforms where a review is facilitated by a different entity than the one where the paper is published (7).

### Open data

Open data are data that can be used (with proper attribution) by anyone without technical or legal restrictions (8). Open Knowledge Foundation characterized them by: a) availability and access:; b) reuse and re-distributionc) universal participation (9). Many statements and recommendations were made to increase data sharing and data citation (10), of which International Committee of Medical Journal Editors (ICMJE) recommendations, followed today by more than 500 biomedical journals, required a data sharing statement for clinical trials from July 2018 (11). Research data is thought to be best preserved by being deposited in one of the numerous general or specific repositories exsisting today (12). Although research and funding agencies often recognize the importance of data sharing, they till face many technical and even psychological barriers to data sharing (13). Nevertheless, there are indications that data sharing has increased with the proliferation of preprint servers and with an increase in pace of scientific communication exchange (14).

### Preprinting

While experiments with faster dissemination of research began in 1960s, in 1990s, first preprint servers (arXiv, SSRN and RePec) emerged and allowed sharing of scholarly manuscripts between researchers before they are peer reviewed. However, it took a while for those servers to become the go to place for researchers. For example, only 8 years after its inception, did arXiv become a major player in the dissemination of results in physics and mathematics (15). Other scholarly fields have been slower to adapt to the preprint culture: with bioRxiv, a preprint server dedicated to the biological sciences, originating in 2013, and MedRxiv, a server for clinical research preprints, in June of 2019 (16). Today, there are more than 60 preprint servers in the world covering all scientific fields (17), and the number of preprints is on the rise, fueled even by the the COVID-19 pandemic (18).

### Attitudes towards open data, preprinting and peer-review

Attitudes influence human intentions and behavior (19). Most research up to now, however, has measured attitudes toward open science using single item questions and analysis of answers on those questions. Although one-question measuring is a useful method for “snapshot measuring” (20), measuring attitudes with only a single question is generally not considered an optimal approach. To the best of our knowledge, no studies, except the Brehm et al study (21) and Curty et al. (22), used a scale (and reported on its reliability and validity) to measure attitudes toward any aspect of open science. It was, therefore, our goal to construct and validate such a scale and use it to explore attitudes of Croatian scientists towards it.

## Methods

A cross-sectional study with psychometrical validation of a questionnaire, which we named, the Attitudes towards Open data sharing, preprinting, and peer-review (ATOPP).

### Participants

In 2018, Croatia had 17,706 scientists (23). In order to reach as most of them as we could, we sent invitations through 2 different channels: through the mailing list of Croatian scientists (approximately 17,000 members) compiled by the Rudjer Boskovic Institute (Zagreb, Croatia), and the Dean’s secretaries of University of Rijeka (the University of the first author, with 1,256 scientists).

### Procedure

Participants were invited to fulfill an anonymous online questionnaire (through Google forms). The survey was open from 12 May 2020 to 7 July 2020, and we sent two reminders 14 days apart.

#### Constructing the questionnaire

The questionnaire that was sent was constructed as a result of three focus groups we held at the University of Rijeka in 2019 and 2020 with a total of 24 participants. The first focus group was held with participants from Biomedical Sceinces (N=12), second with the participants from Social Sciences (N=7) and the last with participants from Natural Sciences (N=5). Participants were asked 5 questions: (1) What is open science to you? (2) What are your experiences with open access journals? (3) What do you think about the open peer-review process? (4) Do you use any of the open science tools? (5) What could influence you to provide access to your research/project data? The sessions were recorded and the transcripts used for generating the survey questions (24). The questionnaire was then facially validated by us (the authors). This sent questionnaire had 72 questions, of which 45 were meant to assess the attitudes towards open science, specifically open access (11 items), open peer-review (15 items), open data (13 items), preprints (14 items), and open science tools (12 items). Then 20 questions inquired on open science practices; and 8 about demographic information. Answers to attitude statements were offered on a five-point Likert-type scale, where 1 indicated “strongly disagree;” 2 – “disagree;” 3 – “neither agree nor disagree;” 4 – “agree;” and 5 – “strongly agree.” Open science practices questions were of mixed type (yes/no and multiple-choice questions). Demographic questions included questions on gender, age, scientific filed, roles in science and number of published papers.

Our initial exploration (factor analysis) of the 45 attitudes questions showed that questions on open access (8 items) and open science tools (6 items) explained less than 5% of the variance of the total score and were not internally consistent (with Cronbach alpha scores <0.65) (25). We then reexamined them (face validity), and hypothesized this is most likely due to the fact that these two aspects of open science dealt with concepts outside of direct researcher’s influence (i.e. they were built by other actors), while data sharing, open peer review, and self-archiving through preprints were under direct (self-)agency of the researchers. The psychometrical validation of the remaining questions (31 items) is presented in the results.

### Statistical analysis

#### Validation of the ATOPP questionnaire

Construct validity of the scale was tested with exploratory factor analysis after the suitability of the item correlation matrix was checked with the Kaiser-Meyer-Olkin (KMO) measure of sampling adequacy and Bartlett’s test of sphericity. In Exploratory Factor Analysis, we used Principal Axis Factoring (PAF) as the factor extraction method and Oblimin as the rotation method. We included the extracted factors with the eigenvalue >1, more than 5% of the construct variance and those which passed visual inspection on the scree plot. Factor loadings <0.30 are not presented (25). The factor analysis procedure uses the pattern of correlation between questionnaire items, which represent directly measured manifest variables, grouping them by the variance they share which is captured by factors that are interpreted as latent dimensions, inferred constructs that are not directly measured. Consequently, each extracted factor or dimension is defined only by questionnaire items to which it relates (25,26). Correlations of factors were calculated with Pearson’s coefficient of correlation.

#### Internal consistency of the scale and subscales were determined with Cronbach alpha

##### Total score

Before calculating the total score we have recorded 4 items: item 6 and 8 in Open data and items 10 and 11 in Open peer-review (Appendix 1). The total score of whole scale and factors were constructed as a linear composite of all items divided by the 21 (number of items) with the score range being from 1 to 5. Lower results (<2.6) were considered as negative attitude, average (2.6-3.39) as neutral attitude and higher results (>3.39) as positive.

#### Analysis of answers, based on the ATOPP survey

Qualitative data are presented with frequency and relative frequency. Comparison of qualitative data is done with χ2 test and test of proportion.

Quantitative data are presented with median and interquartile range (Md(IQR) and the distribution was tested with Kolmogorov-Smirnov test. Comparison of quantitative data was made with non-parametric (Mann-Whitney or Kruskal-Wallis) tests. Post-hoc test for Kruskal-Wallis was Dunn test.

For the purpose of the attitude analysis we have merged Natural sciences and Technical sciences, Biomedicine and health and Biotechnical sciences, and finally Social Sciences, Humanities and Interdisciplinary fields of science.

For statistical analysis, we have used 2 statistical packages SPSS (IBM SPSS Statistics for Windows, Version 25.0. Armonk, NY: IBM Corp) and Medcalc (MedCalc Software, Ostend, Belgium, version 16.0.3). P<0.05 was considered significant.

#### Sample size calculation

It is important to acknowledge that our main goal was construction of the questionnaire, and not the assessment of attitudes of the total or representative population of Croatian scientists. Therefore, we based our calculation on the number of initial survey attitude questions (n=44) and the fact that it is considered sufficient for scale validation to have 10 times more participants than the number of items (27).

##### Ethics

The study was approved by the Ethical committee of the University of Rijeka, Rijeka, Croatia (KLASA: 003-08/19-01/l; URBROJ: 217 0-24-04-3-19-7). In the invite letter we also presented the consent form, which the participants gave by partaking in the survey.

## Results

### Validation of the ATOPP questionnaire

Thirty-one item related to open peer-review (12 items), open data (10 items) and preprinting (9 items) were entered into the exploratory factor analysis after exclusion of 14 items related to open access and open science tools (see methods above). Acceptability of the construct was then assessed by analysing the floor and ceiling effects of the individual items on the score distribution and no floor and ceiling effects were observed.

Kaiser-Mayer Olkin test (KMO=0.79) and the Bartlett’s test of sphericity (P < 0.001), which satisfied the condition for principal axis factoring (PAF) of the 31 item. The inspection of the scree plot, Eigenvalues >1 and more than 5% of variance explained yielded 4 factors that with 40% of the construct variance explained. We then repeated the factor analysis with 22 items that had factor loadings higher than 0.30 (25). The second PAF analysis was more suitable (KMO=0.80; Bartlett’s test P<0.001) and it resulted with 4 factors (21 items) – Open Data, Preprinting, Open peer review in a small scientific communities and Open Peer-review that accounted with 51% of the construct variance (Table 1).

**Table 1.**
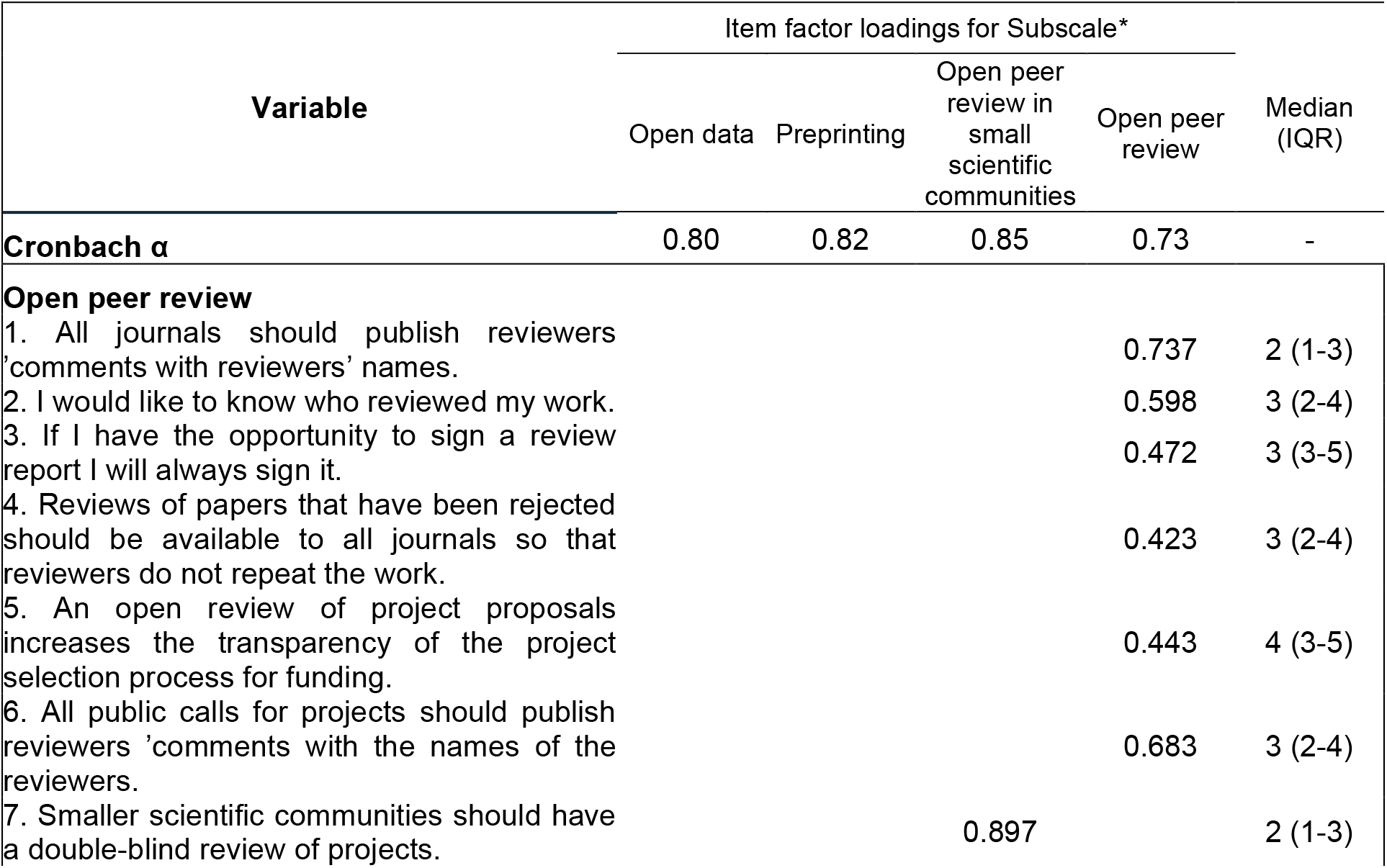
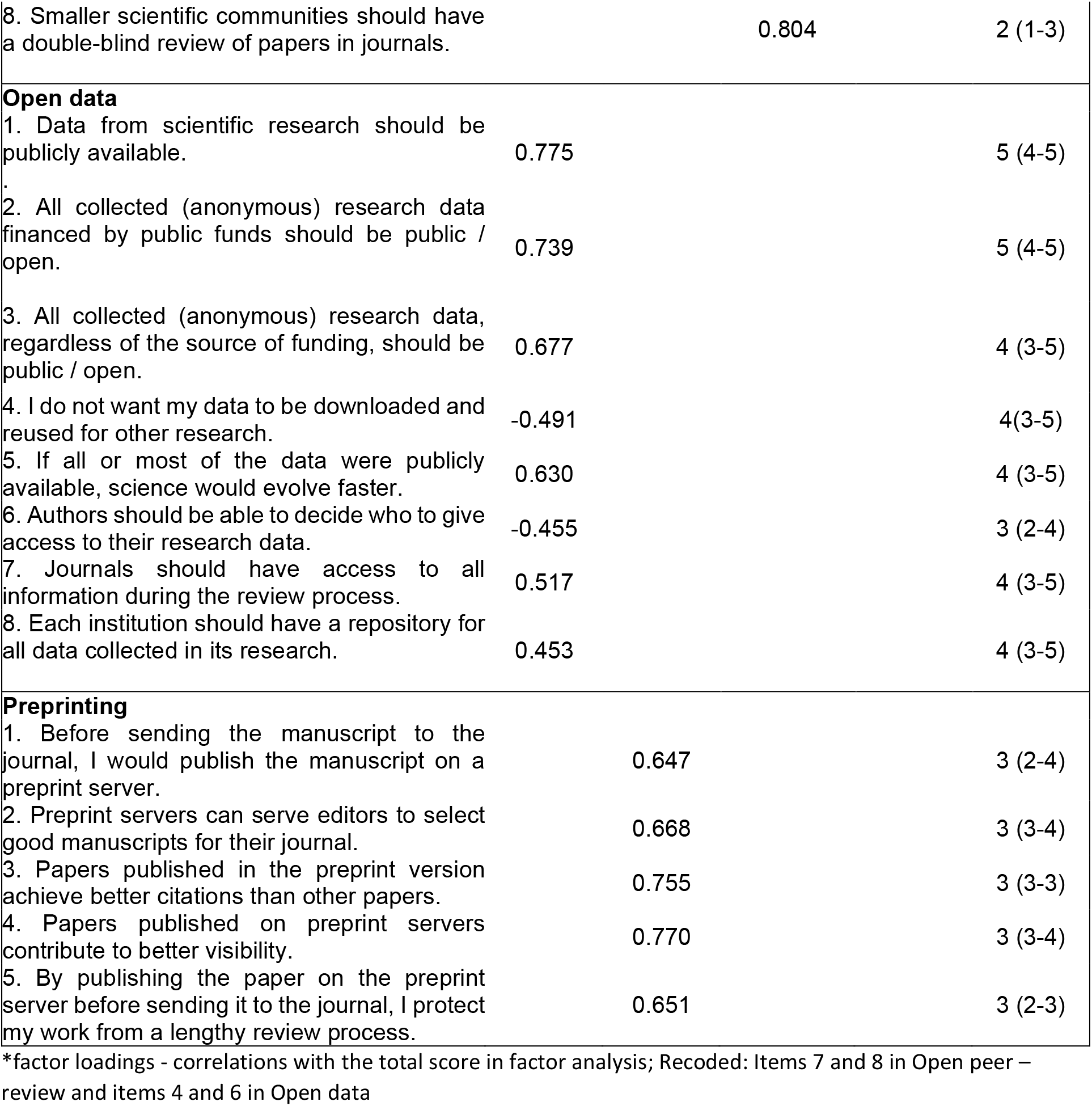
Attitudes towards Open Data, Preprinting, and Peer-review (ATOPP) – reliability, factor loadings and median values.

Structure matrix (correlations of each item with the extracted dimensions) is presented in Appendix 1 Table 1, indicating 21 items were left in the model with a simple factorial structure (loadings are distributed on one factor exclusively). The reliability of the whole scale was very good, Cronbach α= 0.815.

### Participants characteristics

We have collected 546 responses, 196 (36%) from University in Rijeka and 350 (64%) from the Rudjer Boskovic Institute list of Croatian scientists. There was no overlap between the respondents of the two sources, and 5 responses were not valid (not completed), leaving a total of 541 responses. The response rate for the University of Rijeka was 15.6% and for the Croatian scientist list it was 2%. Factorial structure of the attitude scales was the same for both samples and therefore we present them together.

Median age of the participants was 45 (38 to 53), with equal percentage of both males and females (43% vs 54%, P = 0.082) Majority of the respondents were from Biomedicine and Health (26%), Social (25%) or Natural Sciences (17%). They were most commonly Assistant Professors (29%), Full Professors (27%) or Associate Professor (19%). Most respondents (n=529, 98%) published at least one article, with a median of 23 (IQR 10-45). More than two thirds (n=371, 69%) were also reviewers, 16% (n=87) acted as reviewers for funding agencies, and 11% (n=62)as members of the editorial board, finally 3% (n=18) were editors. Detailed demographic and scholarly information of respondents is presented in Table 2.

**Table 2.**
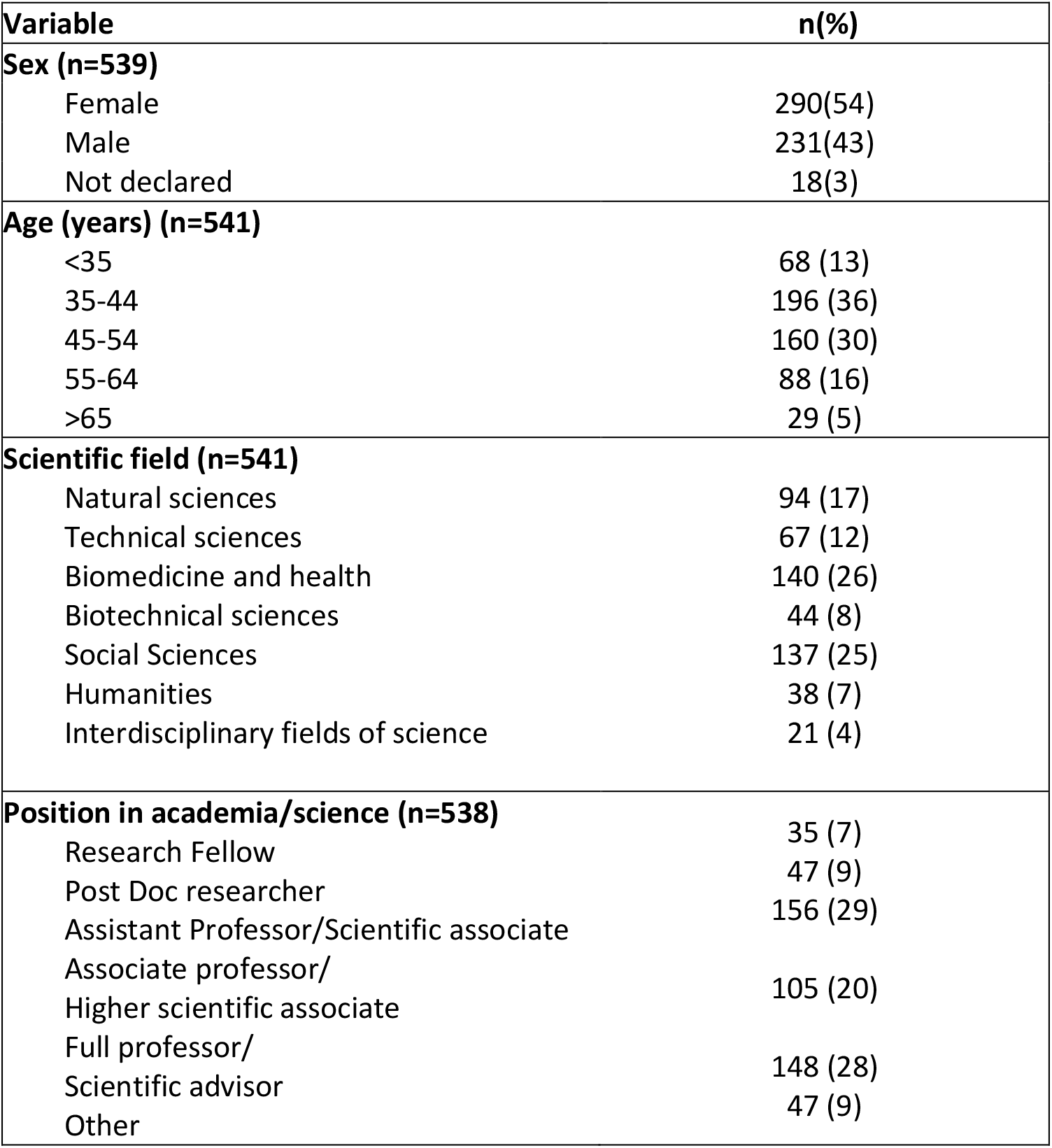
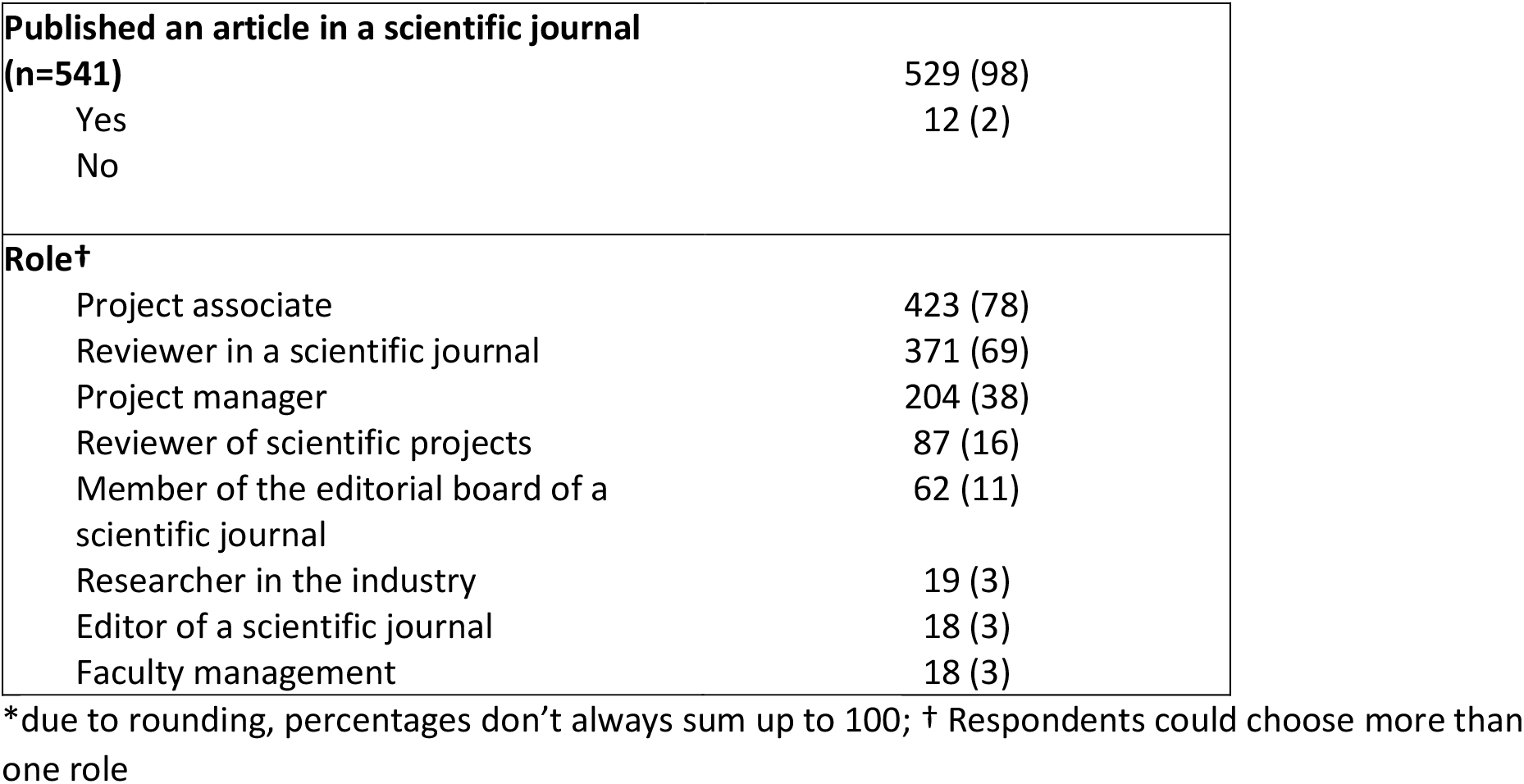
Study participants characteristics (n=541)

### Open science practices

Respondents open science practices are presented in Table 3. Around half (47%, n=240) of the respondents participated in open peer-review and most of them were happy to sign the review reports (n=225, 95%).

**Table 3.**
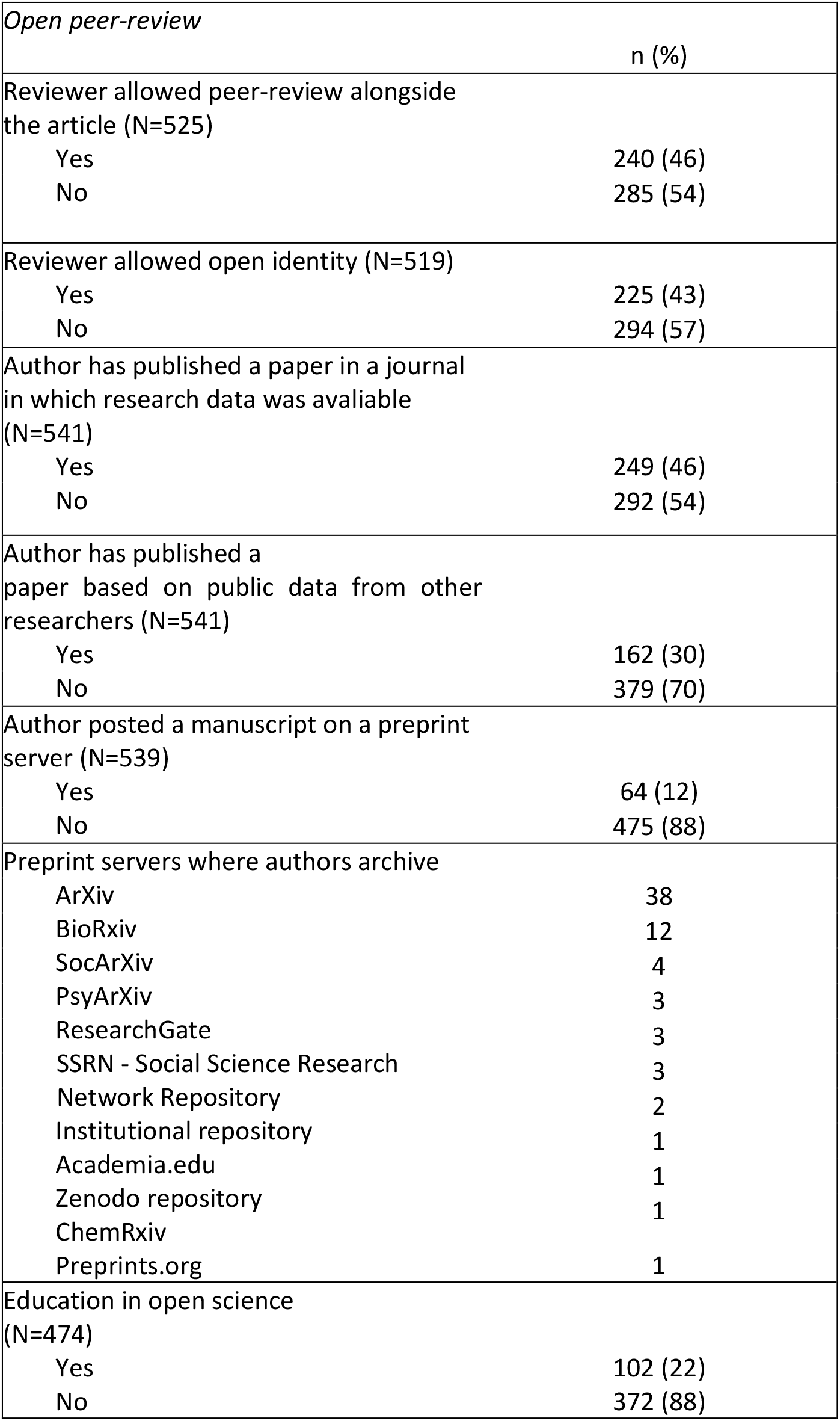
Open peer review, open data and preprinting practices.

| *Open peer-review* |  |
| --- | --- |
|  | n (%) |
| Reviewer allowed peer-review alongside the article (N=525) |  |
| Yes | 240 (46) |
| No | 285 (54) |
| Reviewer allowed open identity (N=519) |  |
| Yes | 225 (43) |
| No | 294 (57) |
| Author has published a paper in a journal in which research data was available (N=541) |  |
| Yes | 249 (46) |
| No | 292 (54) |
| Author has published a paper based on public data from other researchers (N=541) |  |
| Yes | 162 (30) |
| No | 379 (70) |
| Author posted a manuscript on a preprint server (N=539) |  |
| Yes | 64 (12) |
| No | 475 (88) |
| Preprint servers where authors archive |  |
| ArXiv | 38 |
| BioRxiv | 12 |
| SocArXiv | 4 |
| PsyArXiv | 3 |
| ResearchGate | 3 |
| SSRN - Social Science Research | 3 |
| Network Repository | 2 |
| Institutional repository | 1 |
| Academia.edu | 1 |
| Zenodo repository | 1 |
| ChemRxiv | 1 |
| Preprints.org | 1 |
| Education in open science (N=474) |  |
| Yes | 102 (22) |
| No | 372 (88) |

Nearly half of the authors (46%, n=249) published a paper in a journal in which research data could be deposited, and one third (29.9%, n=162) published an article based on public data from other researchers. Most respondents shared their data (as supplementary files) via journals (54%, n=285). Minority of the respondents posted a preprint (12%, n=64), mostly on Arxiv (n=38), BiorXiv (n=12) or SocarXiv (n=4).

### Attitudes towards Open Data, Preprinting, and Peer-review

The total score for all participants on the ATOPP scale was neutral with median of 3.3 (3.0-3.7). The neutral score was also found for their attitudes towards preprinting [3.0 (2.6-3.4)] and open peer review [3.2 (2.7-3.7)]. Negative attitude was found for the open peer-review in small scientific communities [2.0 (1.0-3.0)] and positive for open data [3.9 (3.4-4.4)] (all P<0.05) (Table 4). Differences in attitudes were tested regarding gender, field, open science practices and education (Table 4).

**Table 4.**
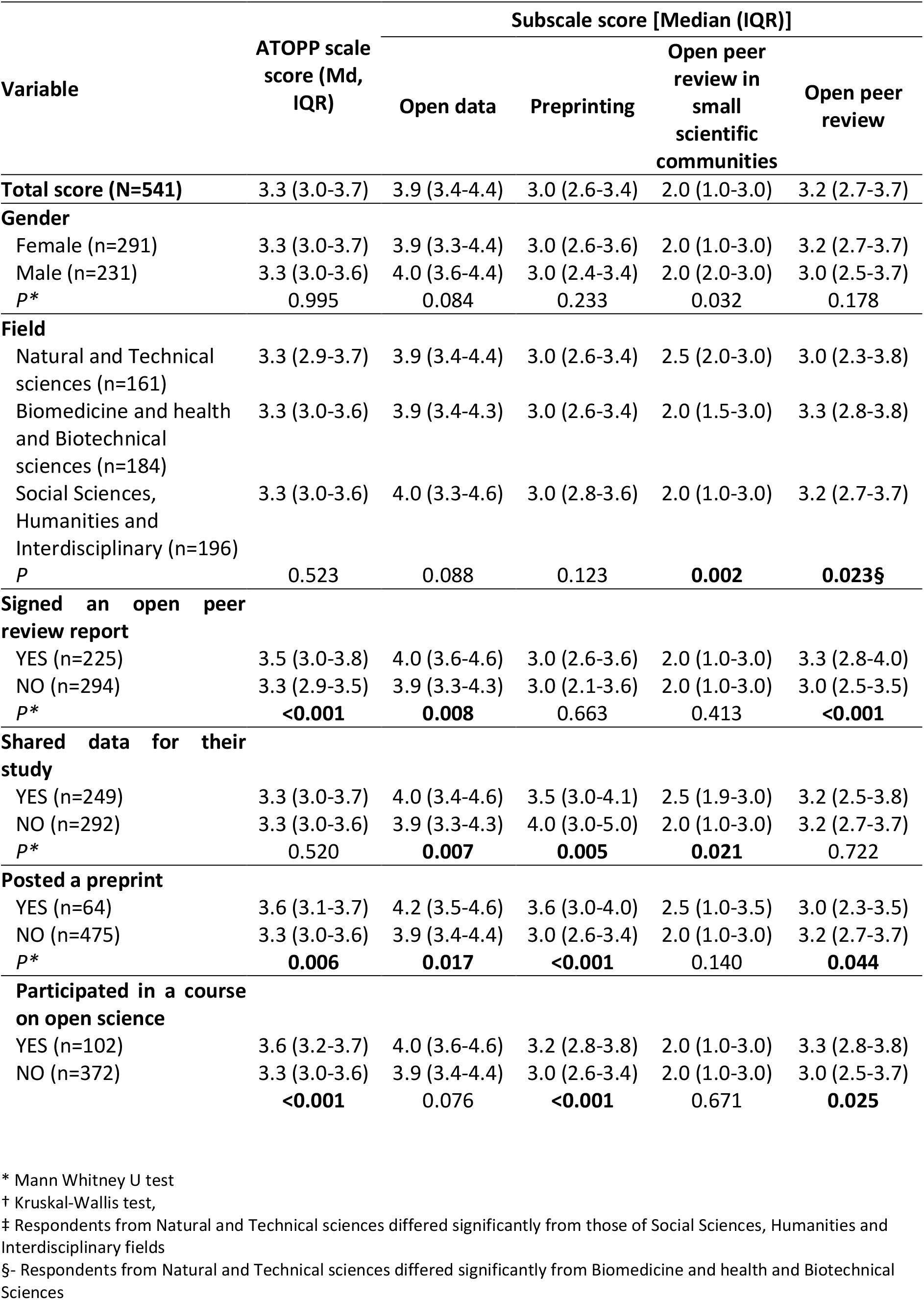
Attitude towards Open Data Sharing, Preprinting, and Peer-review (ATOPP) of Croatian scientists (N=541).

| Variable | ATOPP scale score (Md, IQR) | Subscale score [Median (IQR)] |  |  |  |
| --- | --- | --- | --- | --- | --- |
|  |  | Open data | Preprinting | Open peer review in small scientific communities | Open peer review |
| **Total score (N=541)** | 3.3 (3.0-3.7) | 3.9 (3.4-4.4) | 3.0 (2.6-3.4) | 2.0 (1.0-3.0) | 3.2 (2.7-3.7) |
| **Gender** |  |  |  |  |  |
| Female (n=291) | 3.3 (3.0-3.7) | 3.9 (3.3-4.4) | 3.0 (2.6-3.6) | 2.0 (1.0-3.0) | 3.2 (2.7-3.7) |
| Male (n=231) | 3.3 (3.0-3.6) | 4.0 (3.6-4.4) | 3.0 (2.4-3.4) | 2.0 (2.0-3.0) | 3.0 (2.5-3.7) |
| *p* * | 0.995 | 0.084 | 0.233 | 0.032 | 0.178 |
| **Field** |  |  |  |  |  |
| Natural and Technical sciences (n=161) | 3.3 (2.9-3.7) | 3.9 (3.4-4.4) | 3.0 (2.6-3.4) | 2.5 (2.0-3.0) | 3.0 (2.3-3.8) |
| Biomedicine and health and Biotechnical sciences (n=184) | 3.3 (3.0-3.6) | 3.9 (3.4-4.3) | 3.0 (2.6-3.4) | 2.0 (1.5-3.0) | 3.3 (2.8-3.8) |
| Social Sciences, Humanities and Interdisciplinary (n=196) | 3.3 (3.0-3.6) | 4.0 (3.3-4.6) | 3.0 (2.8-3.6) | 2.0 (1.0-3.0) | 3.2 (2.7-3.7) |
| *P* | 0.523 | 0.088 | 0.123 | **0.002** | **0.023§** |
| **Signed an open peer review report** |  |  |  |  |  |
| YES (n=225) | 3.5 (3.0-3.8) | 4.0 (3.6-4.6) | 3.0 (2.6-3.6) | 2.0 (1.0-3.0) | 3.3 (2.8-4.0) |
| NO (n=294) | 3.3 (2.9-3.5) | 3.9 (3.3-4.3) | 3.0 (2.1-3.6) | 2.0 (1.0-3.0) | 3.0 (2.5-3.5) |
| *p* * | **<0.001** | **0.008** | 0.663 | 0.413 | **<0.001** |
| **Shared data for their study** |  |  |  |  |  |
| YES (n=249) | 3.3 (3.0-3.7) | 4.0 (3.4-4.6) | 3.5 (3.0-4.1) | 2.5 (1.9-3.0) | 3.2 (2.5-3.8) |
| NO (n=292) | 3.3 (3.0-3.6) | 3.9 (3.3-4.3) | 4.0 (3.0-5.0) | 2.0 (1.0-3.0) | 3.2 (2.7-3.7) |
| *p* * | 0.520 | **0.007** | **0.005** | **0.021** | 0.722 |
| **Posted a preprint** |  |  |  |  |  |
| YES (n=64) | 3.6 (3.1-3.7) | 4.2 (3.5-4.6) | 3.6 (3.0-4.0) | 2.5 (1.0-3.5) | 3.0 (2.3-3.5) |
| NO (n=475) | 3.3 (3.0-3.6) | 3.9 (3.4-4.4) | 3.0 (2.6-3.4) | 2.0 (1.0-3.0) | 3.2 (2.7-3.7) |
| *p* * | **0.006** | **0.017** | **<0.001** | 0.140 | **0.044** |
| **Participated in a course on open science** |  |  |  |  |  |
| YES (n=102) | 3.6 (3.2-3.7) | 4.0 (3.6-4.6) | 3.2 (2.8-3.8) | 2.0 (1.0-3.0) | 3.3 (2.8-3.8) |
| NO (n=372) | 3.3 (3.0-3.6) | 3.9 (3.4-4.4) | 3.0 (2.6-3.4) | 2.0 (1.0-3.0) | 3.0 (2.5-3.7) |
|  | **<0.001** | 0.076 | **<0.001** | 0.671 | **0.025** |

\* Mann Whitney U test
† Kruskal-Wallis test,
‡ Respondents from Natural and Technical sciences differed significantly from those of Social Sciences, Humanities and Interdisciplinary fields
§- Respondents from Natural and Technical sciences differed significantly from Biomedicine and health and Biotechnical Sciences

We found no gender differences (all P>0.05) except for the open peer-review in the small scientific communities, where female respondents had a more negative attitude than male respondents [2.0(1.0-3.0) vs 2.0(2.0-3.0), P=0.032].

We also found no differences in the overall ATOPP score between scientific fields (P=0.523). However, attitudes toward open peer review in small scientific communities were higher in Natural sciences and Technical sciences than in Social Sciences, Humanities and Interdisciplinary fields [2.5 (2.0-3.0) vs 2.0 (1.0-3.0), P=0.002]. While attitudes towards open peer review were higher in Biomedicine and Health and Biotechnical sciences compared to Natural sciences and Technical sciences [3.3 (2.8-3.8) vs 3.0 (2.3-3.8), P=0.023)].

Participants who had open peer review experience had higher total ATOPP score (P<0.001), as well as attitudes towards open data (P=0.008) and open peer-review (P<0.001). Similarly, those who previously shared their data had higher attitudes towards Open data (P=0.007), Preprinting (P=0.005) and Open peer review in small scientific communities (P=0.021). Participants with experience in preprinting had more positive attitudes (all P<0.05) except for the Open peer review in small scientific communities (P=0.140). Finally, participants who had education in open science had a more positive ATOPP score then those that did not (<0.001) and they also had higher attitudes except for the open peer-review in small scientific communities.

## Discussion

Our study showed the development of the ATOPP questionnaire for measuring attitudes toward open data, open peer review and preprints. To the best of our knowledge, this is the first psychometrically validated scale for measuring attitudes towards those topics using a multiple-question (scale) approach. The questionnaire demonstrated good internal consistency and construct validity.

During the questionnaire development, attitudes towards open peer review in small scientific communities turned out to be a separate construct (subscale) from attitudes toward open peer review. This could be a product of both the fact that Croatian scientific community for centuries had a higher number of specialized journals per capita compared to its neighboring countries, and the fact that open peer review in small (national) fields or subfields has higher likelihood of reviewers being direct competitors for both funding and job positions (28).

Based on the validated scale, Croatian scientists that answered our survey, had a generally neutral attitude toward open data, open peer review and preprints, and a negative attitude towards open peer review in smaller scientific communities We found no overall gender or field differences. However, scientists who already provided open peer review reports in the past or posted preprints had generally more positive attitudes than those who did not.

It is likely that the positive attitudes towards open data in our study are associated with the high prevalence of researchers in our sample (46%) that shared their data in the past. In recent survey in the United Kingdom with 1724 participants 21% of them have had experience depositing primary data in online repositories although they have very positive opinion about data sharing (13). In a recent survey among members of the German psychological society (N=303) Abele-Brehm et al construct a scale measuring hopes (10 items, Cronbach α=0.90) and fears (4 items, Cronbach α=0.67) towards data sharing, and the positive attitude – “hopes” were neutral (around 3.0), but their experience with data sharing was not measured (21).

Despite studies showing that manuscripts posted first on preprint servers receive more citations than those not posted (16) attitudes towards preprinting were neutral in our study. Although the attitude of participants who had experience with preprinting is positive, it is a small subsample, and only 11% have posted a preprint previously. These self-reported posting is in the line with a large analysis of preprints (2013-2019) on Biorxiv (n=67885), where preprints last author mostly originate from the United States (39.2%) and the United Kingdom (10.5%), while Croatia was described as a “contributor country” – contributor in collaboration, rarely a senior author. Our results are lower than in a recent survey of authors from Latin America (SCIELO) where preliminary results of a survey indicate there were around 40% of authors who posted a preprint and they mostly feel posting a preprint is something beneficial (29). In a recent analysis of comments of 1,983 bioRxiv preprints (before the COVID pandemic) Malicki et al (30) found that 12% (N=168) of comments were full peer review reports. and hypothesised beneficial effects of the comments for the authors and the scientific community.

Attitudes towards open peer review were neutral and lower than expected for the scientist from the Natural sciences and Technical sciences. As expected, scientists who had experience with open peer review had a higher and positive attitude towards open peer review. In a large non psychometrically-validated survey (N=3062) by Ross-Hellauer, Deppe and Schmidt (2017) general attitude towards open peer review was positive, although participants were not keen on signing the review reports, a result we have also obtained with the extraction of the factor Attitude towards open peer review in small scientific communities (31).

This scale, open peer-review, also included questions about project proposals and the need for openness in reviewing scientific projects, which are also important and rarely investigated and one item was retained after the factorial analysis (item: All public calls for projects should publish reviewers ‘comments with the names of the reviewers). Although we have expected scientists would like project proposals review to be open the opinion in our sample was neutral.

Attitude towards open peer review in small scientific communities are low and much lower than attitudes towards open peer-review indicating that Croatian scientific community isn’t ready for this change. Female scientists have a lower attitude than male scientists probably due to well-known reasons of gender balance in academia and fear of vengeance in case of signing a negative review. Scientists who have shared data, have experience in preprinting and from natural and technical sciences have also negative attitude towards peer review in small scientific communities but higher attitude than the rest.

Despite presenting the first psychometrically validated scale for measuring attitudes towards open data, preprinting and open peer-review, our study is not without limitations. As all questionnaires, are data are based on self-declared attitudes and practices and does not capture independently confirmed practices. Furthermore, as many recent online surveys our response rates were low, and this could have also been influenced as the questionnaire was sent during the early months of the pandemic. Furthermore, we might have captured opinions only of those interested in these topics, and the attitudes of Croatian scientists could be even lower. While we did provide definitions of open science practices in our questionnaire, as most of our respondents did not have education in open science, it is possible some held different ideas of those practices.

In conclusion, our study presents one of the first validated questionnaires measuring open science attitudes using a multi-question – scale approach. Further studies are needed to assess attitudes of researchers in other countries, as well as to track changes of these attitudes over time. With increase in open science practices, and more and more funders encouraging or mandating them, we belie that validate tools, such as this one, are needed to assess is implementation of those practices.

## Supporting information

ATOPP questions

ATOPP_data_file

ATOPP_appendix

## Data statement

All data are shared as supplementary files of the manuscript.

## Supplementary files

1. Appendix
2. Data
3. Questionnaire items

## References

1. Şentürk R. Toward an open science and society: multiplex relations in language, religion and society -revisiting Ottoman culture-. İslam Araştırmaları Derg. 2011;(6):93–129.

2. OECD. Making Open Science a Reality OECD MAKING OPEN SCIENCE A REALITY. 2015;(25):1–108. Available from: http://dx.doi.org/10.1787/5jrs2f963zs1-en

3. Vicente-Saez R; Martinez-Fuentes C. Open Science now: A systematic literature review for an integrated definition. J Bus Res. 2018;88:428–36.

4. Brown CT. Living in an Ivory Basement: Stochastic Thoughts on Science, Testing, and Programming. 2016.

5. Tennant J, Agarwal R, Baždarić K, Brassard D, Crick T, Dunleavy D, et al. A tale of two “opens”: intersections between Free and Open Source Software and Open Scholarship [Internet]. 2020. Available from: https://osf.io/preprints/socarxiv/2kxq8/

6. Pontika N, Knoth P, Cancellieri M, Pearce S. Fostering Open Science to Research Using a Taxonomy and an ELearning Portal. In: Proceedings of the 15th International Conference on Knowledge Technologies and Data-Driven Business [Internet]. New York, NY, USA: Association for Computing Machinery; 2015. (i-KNOW ‘15). Available from: https://doi.org/10.1145/2809563.2809571

7. Ross-Hellauer T. What is open peer review? A systematic review [version 2; peer review: 4 approved]. F1000Research. 2017;6(588).

8. OECD. Making Open Science a Reality. 2015;(25).

9. Open Knowledge Foundation. What is Open? [Internet]. 2020. Available from: https://okfn.org/opendata/

10. Lammey R. Data sharing and data citation: Join the movement! Eur Sci Ed. 2019;45(3):58–9.

11. ICMJE data sharing [Internet]. Available from: http://www.icmje.org/recommendations/browse/publishing-and-editorial-issues/clinical-trial-registration.html

12. Nature. Recommended Data Repositories [Internet]. Available from: https://www.nature.com/sdata/policies/repositories

13. Zhu Y. Open-access policy and data-sharing practice in UK academia. J Inf Sci. 2020;46(1):41–52.

14. Berenbaum MR. On Mr. Hyslop’s prediction, content archives, and preprint servers. Proc Natl Acad Sci U S A. 2020;117(17):9131–4.

15. Ginsparg P. Preprint Déjà Vu. EMBO J. 2016;35(24):2620–5.

16. Hoy MB. Rise of the Rxivs: How Preprint Servers are Changing the Publishing Process. Med Ref Serv Q [Internet]. 2020;39(1):84–9. Available from: https://doi.org/10.1080/02763869.2020.1704597

17. Malički, M; Jerončić, A; ter Riet, G; Bouter, LM; Ioannidis, J; Goodman, S; Aalbersberg IjJ. Preprint Servers’ Policies, Submission Requirements, and Transparency inReporting and Research Integrity Recommendations. JAMA - J Am Med Assoc. 2020;324(18):1901–3.

18. Fraser N, Brierley L, Dey G, Polka JK, Pálfy M, Nanni F. Preprinting the COVID-19 pandemic. 2020;

19. Ajzen, I; Fishbein M. The influence of attitudes on behaviour. In: Handbook of attitudes and attitudes change: basic principles. Mahwah (NJ, US): Erlbaum; 2005. p. 173–221.

20. Bowling A. Just one question: If one question works, why ask several? J Epidemiol Community Health. 2005;59(5):342–5.

21. Abele-Brehm AE, Gollwitzer M, Steinberg U, Schönbrodt FD. Attitudes Toward Open Science and Public Data Sharing: A Survey among Members of the German Psychological Society. Soc Psychol (Gott). 2019;50(4):252–60.

22. Curty RG, Crowston K, Specht A, Grant BW, Dalton ED. Attitudes and norms affecting scientists’ data reuse. PLoS One. 2017;12(12):1–22.

23. Croatian Bureau of Statistics. Higher Education (Croatia), 2018 - Statistical Reports. 2018.

24. Breen RL. A Practical Guide to Focus-Group Research A Practical Guide to Focus-Group Research. 2007;8265(2006).

25. Spector P. Summated Rating Scale Construction. Vol. 19, Japanese Society of Biofeedback Research. Sage; 1992. 463–466 p.

26. Costello AB, Osbourne JW. Best practices in exploratory factor analysis: Four recommendations for getting the most from your analysis. Pract Assessment, Res Eval. 2005;10(7):1–9.

27. Costello AB, Osbourne JW. Best practices in exploratory factor analysis: Four recommendations for getting the most from your analysis. Pract Assessment, Res Eval [Internet]. 2005;10(7):1–9. Available from: http://graduate.tuiu.edu/res620sum08/Modules/Module03/FactorAnalysis.pdf

28. Sambunjak D, Ivaniš A, Marušić A, Marušić M. Representation of journals from five neighboring European countries in the Journal Citation Reports. Scientometrics. 2008;76(2):261–71.

29. Funk, Kathryn; Meadows, Alice; Mendonça, Alex; Rieger, Oya; Swaminathan S. Preprint authors optimistic about benefits: preliminary results from the #bioPreprints2020 survey [Internet]. Available from: https://asapbio.org/biopreprints2020-survey-initial-results?fbclid=IwAR07rFL9o43Aj8iBT-s005A-hIPs4Zn61naFPk9eJ5nTyf9-8Mk7SZ_rCxo

30. Malički M, Costello J, Alperin JP, Maggio LA. From amazing work to I beg to differ - analysis of bioRxiv preprints that received one public comment till September 2019. 2020;(September 2019):1–16.

31. Ross-Hellauer T, Deppe A, Schmidt B. Survey on open peer review: Attitudes and experience amongst editors, authors and reviewers. PLoS One. 2017;12(12):1–28.

