## Supplementary material for "Attitudes and Practices of Open Data, Preprinting, and Peer-review - a Cross Sectional Study on Croatian Scientists": ATOPP questions

### ATOPP questionnaire questions

#### Open peer review
1. All journals should publish reviewers' comments with reviewers' names.
2. I would like to know who reviewed my work.
3. If I have the option to sign a review report, I will always sign it.
4. Reviews of rejected manuscripts should be available to all journals to prevent reviewers repeating the same work.
5. Open review for project proposals increases the transparency of funding allocation procedures.
6. All public calls for projects proposals should publish reviewers' comments with reviewers' names.
7. Small scientific communities should have double-blind reviews for projects proposals.
8. Small scientific communities should have double-blind reviews for journal papers.

#### Open data
1. Data from scientific research should be publicly available.
2. All (anonymized) research data of publicly funded research should be public/open.
3. All (anonymized) research data, regardless of who funded the research, should be public/open.
4. I do not want my data to be downloadable and reusable in other research.
5. If all or most of data were publicly available, science would develop faster.
6. Authors should be able to decide whom to give access to for their research data.
7. Journals should have access to all data during the review process.
8. Each institution should have a repository for all research data it collects.

#### Preprinting
1. Before sending a manuscript to a journal, I would publish the manuscript on a preprint server.
2. Preprint servers can help editors select good manuscripts for their journal.
3. Papers deposited as preprint versions receive more citations than other papers.
4. Papers deposited on preprint servers help increase visibility.
5. By depositing a paper on a preprint server before submitting it to a journal, I protect my research from a lengthy review process.

#### Practices:

Have you ever allowed publication of your review? Yes/No
If yes, did you sign it?
Consequences of signing your review were:
Positive

Negative
There were no consequences

Have you ever used publicly available data of other researchers and used them as basis of your publication?  Yes/No
I published a paper for which even research data was made available. Yes/No
I deposited my research data in: The journal of publication, Dabar repository, Zenodo repository, Other

I archive my papers: in institution's digital repository, in subject area digital repository, in digital repositories (e.g. Zenodo), on personal website(s), on social netoworks (e.g. ResearchGate, Academia.edu, etc.)

I archive papers: Personally, Others do it in my name: librarians, assistants, administrative personell, etc.

Which version of the paper do you deposit: Manuscript version preceding submission for publication, Manuscript accepted for publication, Published manuscript at the moment of its publication, Published manuscript after several months of it being published in a journal, according to copyright agreement, All versions, Other:

Until now, have you deposited a manuscript on any of the preprint servers? Yes/No
If yes, on which one: arXiv, bioArxiv, socarXiv, Other:

Demographic and publication data:

Please indicate your gender: Female, Male, Prefer not to say

Age (years):

Your area: 1. Natural sciences, 2. Technical sciences, 3. Biomedicine and Health, 4. biotechnical sciences, 5. Social sciences, 6. Humanities, 7. Arts, 8. Interdisciplinary sciences, 9. Interdisciplinary arts

Your current status: Research fellow or assistant; Postdoc; Assistant professor; Associate professor; Professor or above; Other

Which scholarly and research activities do you perform: Project member, Principal investigator, Industry researcher, Reviewer, Editor of a jornal, Editorial board member, Ethics board member, Faculty board member, Other

If you teach, how many teaching hours per year do you teach?

How many scholarly works have you published:

Have you had any open science education until now? Yes/No/Not sure
