## Supplementary material for "Attitudes and Practices of Open Data, Preprinting, and Peer-review - a Cross Sectional Study on Croatian Scientists": ATOPP_appendix

**Appendix 1**

**Factor analysis of the Attitudes and Practices of Open Data, Preprinting, and Peer-review - a Cross Sectional Study on Croatian Scientists**

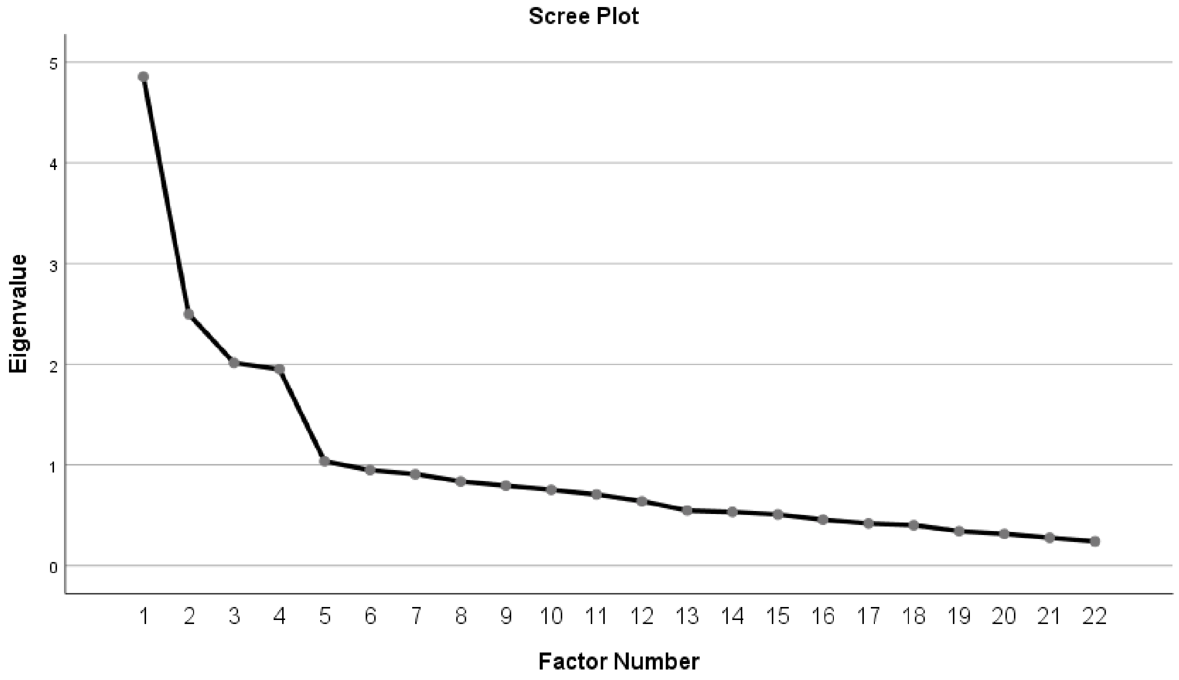

**Figure 1. Scree plot of the factor analysis (22 items) of Attitudes towards Open Data, Preprinting, and Peer-review (ATOPP)**

Factor 1 contained 8 items (1, 2, 3,6,7,8,9,10) and we named it Open data. The reliability of this factor was good, Cronbach α=0.80. Factor 2 contains 5 items (1, 2, 5,6,7) and was named Preprinting. The reliability of this factor was also good, Cronbach α=0.82. Factor 3 contained 2 items (10,11) and was named Open peer review in small scientific communities. The reliability of this scale was also good, Cronbach α=0.85. Factor 4 contains 6 items (1,4,5,6,7,8) and was named Open peer review according to the items meaning. The reliability of this factor was acceptable, Cronbach α=0.73.

The first factor - Open data was positively correlated with other factors (r=0.29 with Factor 2, 0.16 with Factor 3 and 0.34 with Factor 4). The second factor was positively correlated with other factors (r=0.15 with Factor 3 and r=0.35 with Factor 4) and the third and the fourth factor were correlated 0.16.

**Table 1. Attitudes towards Open Data, Preprinting, and Peer-review (ATOPP) –item factor loadings and reliability**

| Variable | Item factor loadings for Subscale ** | | | |
|---|---|---|---|---|
| | Open data | Preprinting | Open peer review in a small scientific community | Open peer review |
| **Cronbach α** | 0.80 | 0.82 | 0.85 | 0.73 |
| **Open peer review** | | | | |
| 1. All journals should publish reviewers 'comments with reviewers' names. | | | | 0.737 |
| *2. All journals should publish reviewers 'comments, but without reviewers' names. | | | | |
| *3. Open review is difficult in smaller scientific communities. | | | | |
| 4. I would like to know who reviewed my work. | | | | 0.598 |
| 5. If I have the opportunity to sign a review report I will always sign it. | | | | 0.472 |
| 6. Reviews of rejected manuscripts should be available to all journals to prevent reviewers repeating the same work. | | | | 0.423 |
| 7. An open review of project proposals increases the transparency of the project selection process for funding. | | | | 0.443 |
| 8. Open review for project proposals increases the transparency of funding allocation procedures. | | | | 0.683 |
| *9. All public calls for projects proposals should publish reviewers' comments with reviewers' names. | | | | |
| 10. Small scientific communities should have double-blind reviews for projects proposals. | | | 0.897 | |
| 11. Small scientific communities should have double-blind reviews for journal papers. | | | 0.804 | |
| *12. Young scientists do not want to sign an open review because they are afraid of the reactions of older colleagues. | | | | |
| **Open data** | | | | |
| 1. Data from scientific research should be publicly available. | 0.775 | | | |
| 2. All (anonymized) research data of publicly funded research should be public/open. | 0.739 | | | |
| 3. All (anonymized) research data, regardless of who funded the research, should be public/open. | 0.677 | | | |
| *4. All data collected in surveys should be available on request. | | | | |
| *5. Research results should be available only to members of the academic community. | | | | |
| 6. I do not want my data to be downloadable and reusable in other research. | -0.491 | | | |
| 7. If all or most of the data were publicly available, science would develop faster. | 0.630 | | | |
| 8. Authors should be able to decide whom to give access to for their research data. | -0.455 | | | |
| 9. Journals should have access to all information during the review process. | 0.517 | | | |
| 10. Each institution should have a repository for all research data it collects. | 0.453 | | | |
| **Preprinting** | | | | |

| | |
|---|---|
| 1. Before sending a manuscript to a journal, I would publish the manuscript on a preprint server. | 0.647 |
| 2. Preprint servers can help editors select good manuscripts for their journal. | 0.668 |
| *3. The publisher / magazine has the right to refuse to publish the work that I previously published on the preprint server. | |
| *4. I do not want to publish a manuscript of the paper before sending it to the journal for fear of stealing the idea, research. | |
| 5. Papers deposited as preprint versions receive more citations than other papers. | 0.755 |
| 6. Papers deposited on preprint servers help increase visibility. | 0.770 |
| 7. By depositing a paper on a preprint server before submitting it to a journal, I protect my research from a lengthy review process. | 0.651 |
| *8. It is enough for me to publish the work on the preprint server. | |
| *9. Before sending the paper to the journal, I check the conditions of copyright and self-archiving. | |

*items removed from the final version of the questionnaire; **the values presented are correlations with the total score; **the values presented are correlations with the total score; Items 4 and 6 in the Open data subscale and Items 10 and 11 in Open peer-review were recoded after the factor anaylsis
